# Refining the Y chromosome phylogeny with southern African sequences

**DOI:** 10.1101/034983

**Authors:** Chiara Barbieri, Alexander Hübner, Enrico Macholdt, Shengyu Ni, Sebastian Lippold, Roland Schröder, Sununguko Wata Mpoloka, Josephine Purps, Lutz Roewer, Mark Stoneking, Brigitte Pakendorf

## Abstract

The recent availability of large-scale sequence data for the human Y chromosome has revolutionized analyses of and insights gained from this non-recombining, paternally inherited chromosome. However, the studies to date focus on Eurasian variation, and hence the diversity of early-diverging branches found in Africa has not been adequately documented. Here we analyze over 900 kb of Y chromosome sequence obtained from 547 individuals from southern African Khoisan and Bantu-speaking populations, identifying 232 new sequences from basal haplogroups A and B. We find new branches within haplogroups A2 and A3b1 and suggest that the prehistory of haplogroup B2a is more complex than previously suspected; this haplogroup is likely to have existed in Khoisan groups before the arrival of Bantu-speakers, who brought additional B2a lineages to southern Africa. Furthermore, we estimate older dates than obtained previously for both the A2-T node within the human Y chromosome phylogeny and for some individual haplogroups. Finally, there is pronounced variation in branch length between major haplogroups; haplogroups associated with Bantu-speakers have significantly longer branches. This likely reflects a combination of biases in the SNP calling process and demographic factors, such as an older average paternal age (hence a higher mutation rate), a higher effective population size, and/or a stronger effect of population expansion for Bantu-speakers than for Khoisan groups.

## MANUSCRIPT

### INTRODUCTION

The Y chromosome phylogeny has been radically revised in the past few years with the advent of next-generation sequencing methods, which revealed thousands of new polymorphic sites (Cruciani et al. 2011; Mendez et al. 2013; Francalacci et al. 2013; Wei et al. 2013; Hallast et al. 2015; Scozzari et al. 2014; Lippold et al. 2014; Karmin et al. 2015). However, the most comprehensive studies were mainly centered on Eurasian samples (Wei et al. 2013; Hallast et al. 2015; Lippold et al. 2014; Karmin et al. 2015). The available sequences to date therefore heavily underrepresent African populations and the haplogroups at the root of the phylogeny, namely haplogroup A and haplogroup B: in total, only 24 sequences from haplogroup A and 46 sequences from haplogroup B were included in the studies cited above. These early-diverging haplogroups comprise sub-branches that are characteristic of different populations and different regions of the African continent (Batini et al. 2011; Scozzari et al. 2014). One of the largest studies of the variation in the basal Y-chromosomal haplogroups A and B published to date, which is based on single nucleotide polymorphisms (SNPs) and short tandem repeats (STRs), localizes these haplogroups largely in Central, East, and Southern Africa, with different subhaplogroups found in each of the geographic regions (Batini et al. 2011). In Southern Africa, three lineages have to date been described as characteristic of the autochthonous populations of foragers and pastoralists, also known as “Khoisan” (Underhill et al. 2000; Wood et al. 2005; Soodyall et al. 2008; Batini et al. 2011). In the nomenclature of the YCC refined in Karafet et al. (2008), which we follow here, these haplogroups are A2, A3b1 and B2b. While haplogroups A2 and A3b1 are restricted to southern Africa, haplogroup B2b is also very frequent in foragers of the Central African rainforest, albeit represented by separate subhaplogroups.

We here use the term “Khoisan” to label populations speaking indigenous non-Bantu languages of southern Africa that make heavy use of click consonants (Güldemann 2014), irrespective of the fact that these populations are linguistically, culturally, and biologically heterogeneous. Similarly, we use the term “Bantu” to refer to the language family that is diffused over vast areas of sub-Saharan Africa (Williamson and Blench 2000), without any racial connotation. Southern African Khoisan groups are known to harbor a remarkable level of genetic variability both for autosomal loci (Pickrell et al. 2012; Schlebusch et al. 2013) and mtDNA sequences (Barbieri et al. 2014a, 2013). However, very little is known about the Y-chromosomal variation in Khoisan groups, as previous studies included data from only a few such populations (Wood et al. 2005; Soodyall et al. 2008; Henn et al. 2011), missing most of the cultural and linguistic diversity subsumed under this generic label (Barnard 1992; Güldemann 2014). Thus, an in-depth Y chromosome study of large numbers of Khoisan individuals is expected to considerably refine our knowledge of the diversity found in the early-diverging haplogroups A and B.

In this study, we use the array designed by Lippold et al. (2014) to generate ~900 kb of Y chromosome sequence data, including off-target variants from the regions flanking the captured SNPs. We apply this method to a dataset of 547 southern African individuals speaking Khoisan and Bantu languages, covering most of the cultural and linguistic diversity of the region (Supplemental Figure S1). Our results reveal new branches within the phylogeny as well as older ages for most of the haplogroups and allow us to reassess previous proposals concerning the diversity and distribution of the early-diverging haplogroups.

### RESULTS

We sequenced ~964 kb of the Y-chromosome from 547 individuals speaking Khoisan and Bantu languages (see Methods). To improve the accuracy of our phylogenetic reconstruction (i.e. to avoid discarding informative positions because they contain missing data), we applied a conservative imputation method: this allowed us to recover a total of 2837 SNPs. As shown in Supplemental Figure S2, the imputation method used here is very robust: even when 50% of the sites are imputed, the error rate is less than 1%. The impact of imputation on the loss of diversity is also minimal, as shown by analyses of a simulated dataset and in subsets of the data that are less-imputed: at most 0.4% of the sites of the entire alignment are affected by a loss of polymorphisms (Supplemental Figure S3A). For an upper boundary of ≤ 10% missing data, no doubletons or tripletons become invariant in the simulations, only singletons (Supplemental Figure S3B).

#### Major southern African haplogroups

The major haplogroups found in our dataset are A2, A3b1, B2a, B2b, and E (including E1a1a, E1a1b and E2); furthermore, individual sequences belonging to haplogroups G, I, O, T, and R1 were found. The phylogeny reconstructed with a Maximum Parsimony tree (Figure 1) and verified by means of network analysis (Supplemental Figure S4-S7) corresponds to that of the ISOGG consortium (International Society of Genetic Genealogy 2014, Version: 10.101, Date: 8 December 2015), as summarized in van Oven et al (Van Oven et al. 2014); however, we identify additional branches that have not yet been reported. Supplemental Table S1 summarizes information about the major branches reported in Figure 1, such as the different nomenclatures used and the mutations defining each branch. The haplogroup assignment for each individual is listed in Supplemental Table S2, while haplogroup frequencies and measures of diversity are shown in Table 1.

**Figure 1.**
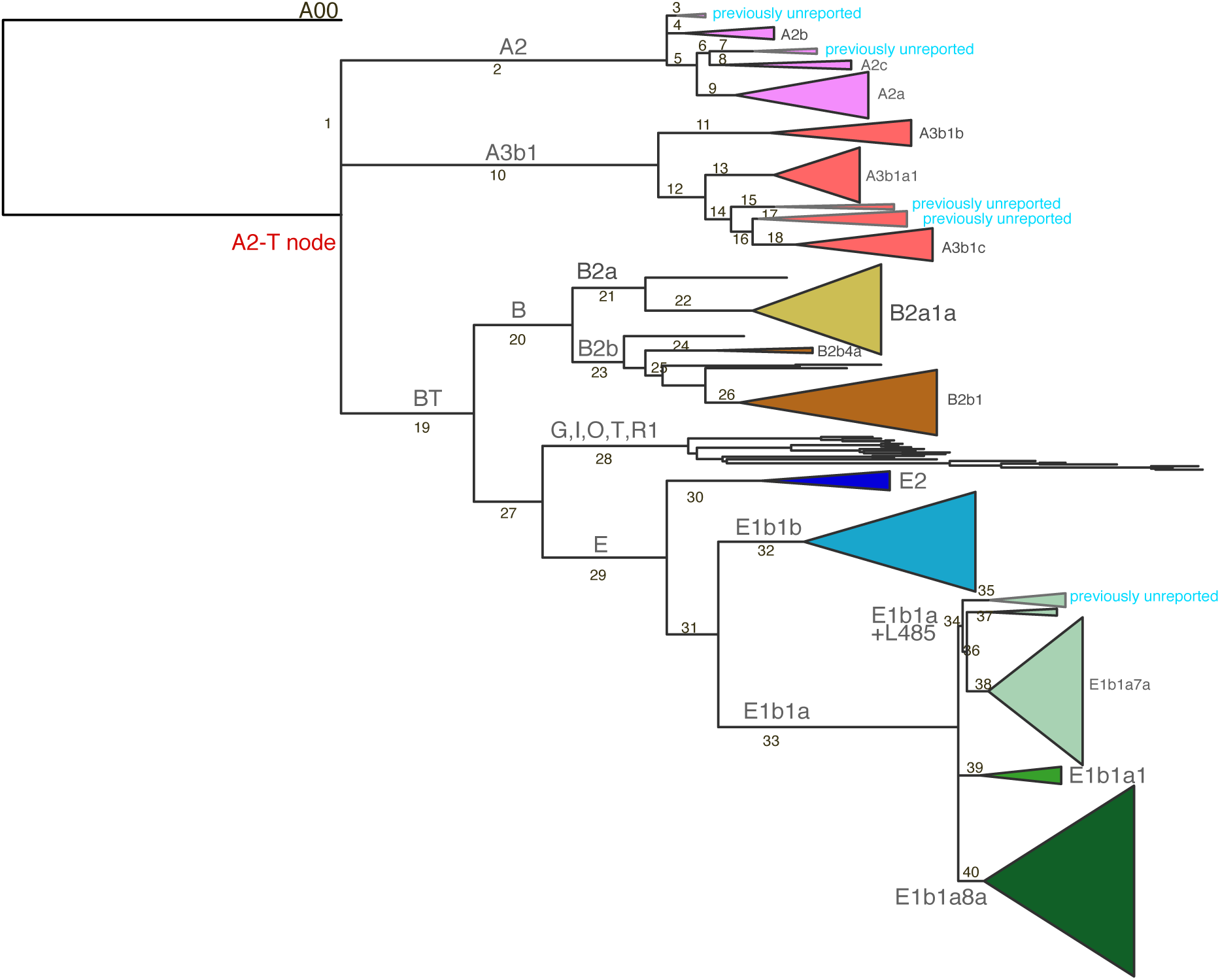
Maximum Parsimony (MP) tree for the southern African dataset, rooted with A00. The width of the triangles is proportional to the number of individuals included. Previously unreported lineages are highlighted. Branches are numbered to identify them in Supplemental Table S1, where information on the defining mutations and comparison with other nomenclature systems are reported. Branch number 1 indicates the branch shared by A2 and A3b1, which is not visible as a separate branch in the MP reconstruction.

**Table 1:** Diversity and other statistics for the major haplogroup branches.

| Major Haplogroup Branch | Sample size | Frequency | Nucleotide diversity | Variance | No. of haplotypes | Haplotype Diversity | SD | Frequency in Khoisan | Frequency in Bantu | p-value |
| --- | --- | --- | --- | --- | --- | --- | --- | --- | --- | --- |
| **A2** | 49 | 0.09 | 0.009 | 0.00002 | 44 | 0.991 | 0.002 | 13.2 | 0.0 | 0.000 |
| **A3b1** | 83 | 0.15 | 0.018 | 0.00007 | 72 | 0.992 | 0.001 | 20.2 | 3.6 | 0.000 |
| **B2a** | 53 | 0.1 | 0.005 | 0.00001 | 38 | 0.913 | 0.015 | 9.2 | 13.6 | 0.195 |
| **B2b** | 47 | 0.09 | 0.013 | 0.00004 | 40 | 0.992 | 0.001 | 11.6 | 2.1 | 0.002 |
| **G,I,O,T,R1** | 20 | 0.04 | 0.042 | 0.00044 | 20 | 1 | 0 | 3.2 | 2.1 | 0.720 |
| **E2** | 12 | 0.02 | 0.005 | 0.00001 | 10 | 0.955 | 0.019 | 1.1 | 5.0 | 0.017 |
| **E1b1b** | 59 | 0.11 | 0.005 | 0.00001 | 47 | 0.978 | 0.003 | 15.1 | 2.1 | 0.000 |
| **E1b1a+L485** | 101 | 0.18 | 0.006 | 0.00001 | 91 | 0.996 | 0.0004 | 8.9 | 35.7 | 0.000 |
| **E1b1a1** | 11 | 0.02 | 0.006 | 0.00001 | 11 | 1 | 0 | 1.1 | 3.6 | 0.125 |
| **E1b1a8a** | 112 | 0.2 | 0.004 | 0.000005 | 91 | 0.966 | 0.004 | 16.4 | 32.1 | 0.000 |

Haplogroup A2, which is defined by 72 mutations (see Supplemental Table S1 for a list of these), includes five monophyletic branches in our data (Supplemental Figure S5), of which only three (A2a, A2b, and A2c) were previously identified in the literature. Of these, A2a is the most frequent.

Haplogroup A3b1 is the only subhaplogroup of A3 present in our dataset, as found previously for southern Africa (Batini et al. 2011). It is the most frequent early-diverging lineage found in our study and is characterized by the highest nucleotide diversity among the major African haplogroups (Table 1). All the individuals within this lineage harbor the defining mutation M51, whereas the P71 mutation is derived only in a subbranch (A3b1a). This agrees with the phylogeny presented previously (Karafet et al. 2008), but contradicts the ISOGG tree, which reports P71 and M51 at the same branching level. We also confirm the diagnostic positions for haplogroups A3b1b (V37) and A3b1c (V306) as defined previously (Scozzari et al. 2012), which are not included in the ISOGG list. However, the three lineages do not split in parallel as reported (Scozzari et al. 2012); rather, A3b1b branches first. Furthermore, we identify two previously undetected clades in between A3b1a1 and A3b1c (Figure 1, Supplemental Figure S5).

B2b and B2a differ notably in their branching structure, as visible from the network (Supplemental Figure S6): B2b exhibits dispersed sequences separated by long branches, while B2a shows a clear star-like expansion, with branches of variable length radiating from a core haplotype. Haplogroup B2b is also commonly found in forager populations of the Central African rainforest (Berniell-Lee et al. 2009; Batini et al. 2011). Here we identify the two branches B2b1 and B2b4a already reported in southern African populations (Batini et al. 2011; Wood et al. 2005), plus four sequences that do not fall in previously reported branches (Supplemental Figure S6).

Whereas most African haplogroups differ significantly in frequency between the Khoisan and Bantu-speaking groups in our study, thereby showing a signature of having a Khoisan vs. Bantu origin in southern Africa, haplogroup B2a does not (Table 1). Moreover, haplogroup B2a is characterized by long branches radiating from a core haplotype found in both Khoisan and Bantu speakers (Figure 2A). As shown by the map in Figure 2B, which visualizes frequency data from these and other African populations (Supplemental Table S3), this haplogroup is widespread over the continent, with the highest frequencies found in populations from Botswana as well as from Cameroon. From these data, it is not clear if haplogroup B2a is an autochthonous Khoisan haplogroup, or a haplogroup brought to southern Africa by Bantu-speakers, or both. To further investigate this haplogroup, we generated STR haplotypes based on 16 loci and compared these to published data; the network generated from these STR haplotypes (Supplemental Table S4, Supplemental Figure S8) shows haplotypes of southern African Khoisan and Bantu speakers located towards the core, and two separate clusters of haplotypes from central Africa and elsewhere at the periphery. Hence, the STR data also do not provide a clear signal of the origin of this haplogroup.

**Figure 2.**
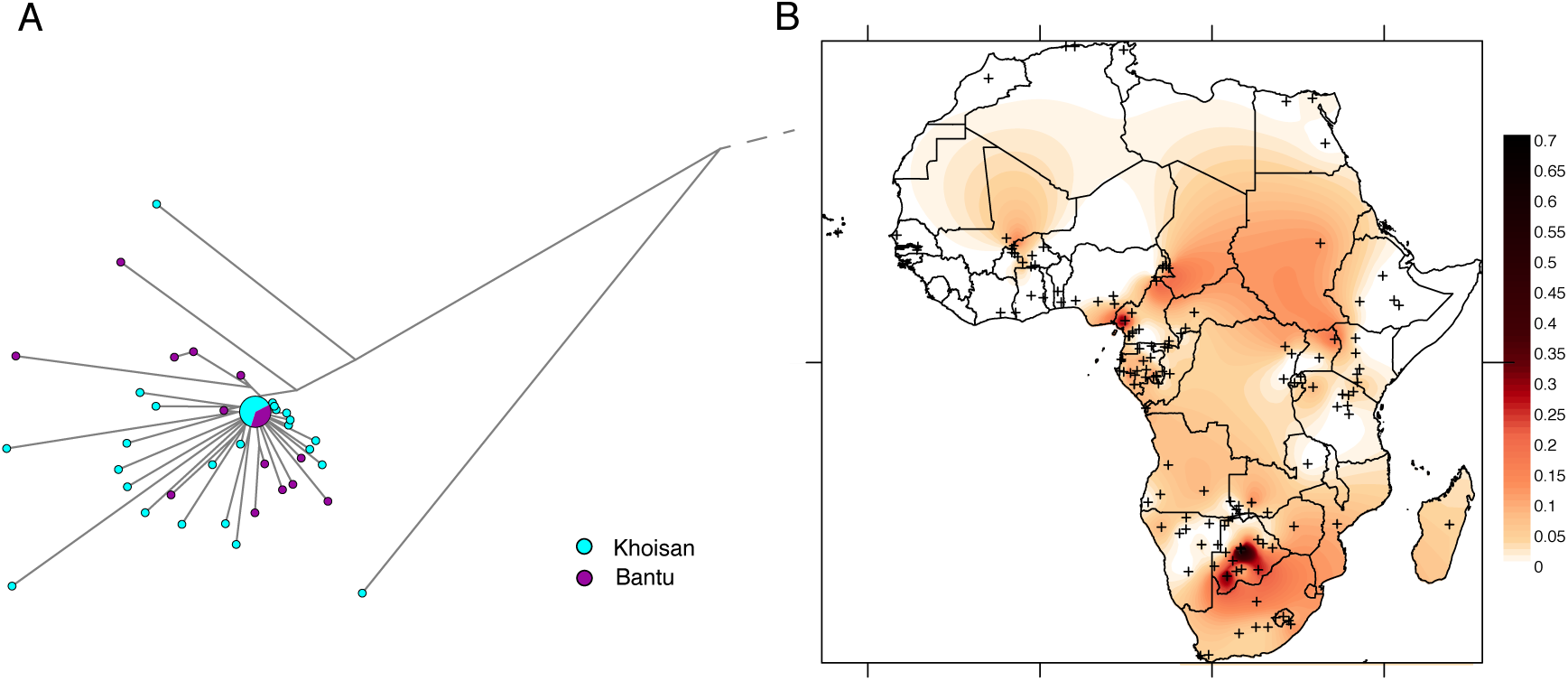
Diversity and distribution of haplogroup B2a. A. Network of B2a sequences color-coded by linguistic affiliation (Khoisan vs. Bantu speaking individuals). The dashed line indicates the position of branch 21 from Figure 1, which leads to the root of B2a. B. Schematic distribution of haplogroup B2a in Africa: the more intense the color, the higher the frequency in the population. Small crosses mark the locations of the 146 African populations included in the analysis (see Supplemental Table S3).

Lastly, within haplogroup E we find E2, E1b1b, and three subgroups of E1b1a, namely E1b1a1, E1b1a8a, and a subgroup characterized by mutation L458, which includes E1b1a7, but which was not recognized previously (Karafet et al. 2008). We here refer to this subgroup as E1b1a+L458 (Supplemental Figure S7).

#### TMRCA and variation in branch length

Estimates of the Time to the Most Recent Common Ancestor (TMRCA) were obtained with two different methods: count of mutations (corresponding to the rho statistic) and BEAST analysis. The TMRCA for the deepest node found in our dataset (A2-T) is 218 kyrs based on counting mutations and 248 kyrs based on BEAST analyses (Figure 3). As shown by the comparison with TMRCA estimates for various nodes obtained by other studies (Figure 3), our estimates are always older than those published previously.

**Figure 3.**
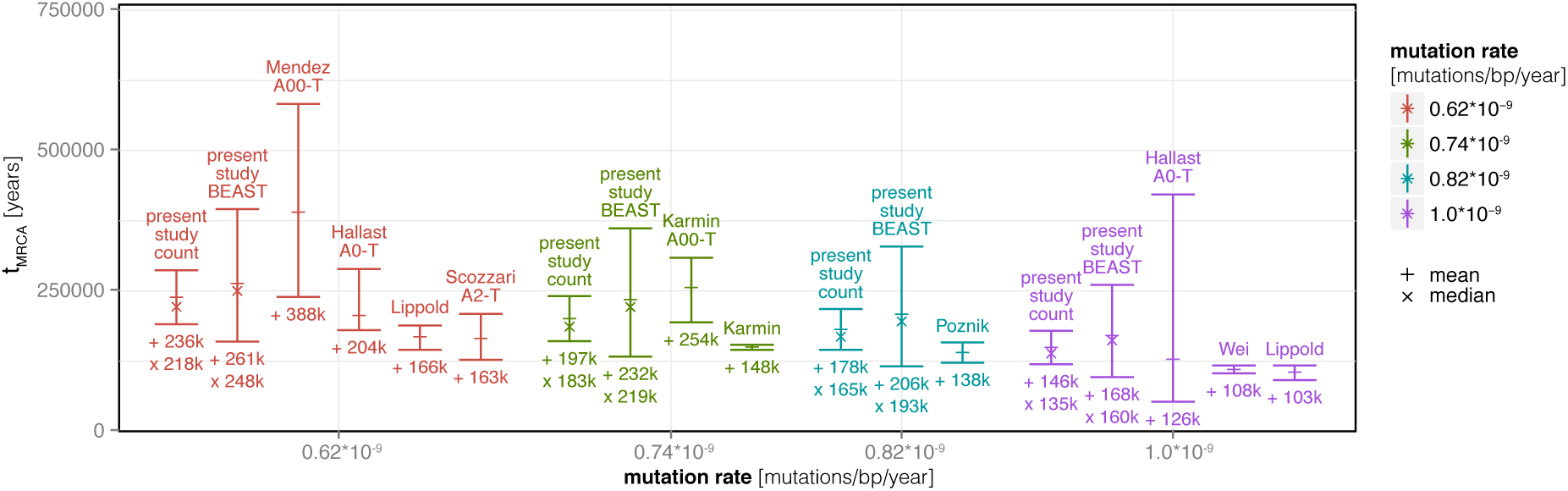
Values of TMRCA for the A2-T node from the present study, estimated by direct count and by BEAST analysis, for four different mutation rates (indicated with different colors); both median and mean estimates are indicated. The dates are compared with estimates from other studies (indicated by the name of the first author), which variously dated the same A2-T node (not explicitly labeled in the figure) or the A00-T or A0-T nodes (identified above the bars).

We also estimated TMRCAs for the individual haplogroups within A and B with three different methods, including calculations based on the count of mutations from the root (Supplemental Table S5), BEAST estimates from the whole phylogeny (Supplemental Figure S9) and independent BEAST estimates from runs for single major haplogroups (Supplemental Figure S10). The dates we obtain are again substantially older than those proposed in the literature, which are based on eight STR loci (Batini et al. 2011). The coalescence of A2 dates to between 27 and 33 kya instead of 6 kya, that of A3b1 to 47–64 kya instead of 10 kya, that of B2b is dated to 46–74 kya, and that of B2a to 46–51 kya (Supplemental Figure S10). Bayesian Skyline Plots (BSPs) computed for the major haplogroups all display population expansions of varying degrees coinciding with the beginning of the Holocene, ~7–12 kya (Supplemental Figure S11).

An analysis of the distribution of the number of mutations from each tip to the A2-T node (Figure 4A) demonstrates considerable heterogeneity in branch length, with a bimodal distribution. Furthermore, the branch lengths differ strikingly among haplogroups (Figure 4B): A, B, E2 and E1b1b are characterized by shorter than average branch lengths, while the E1b1a subgroups all have significantly longer branches (Wilcoxon test W = 71048, p-value < 0.001).

**Figure 4.**
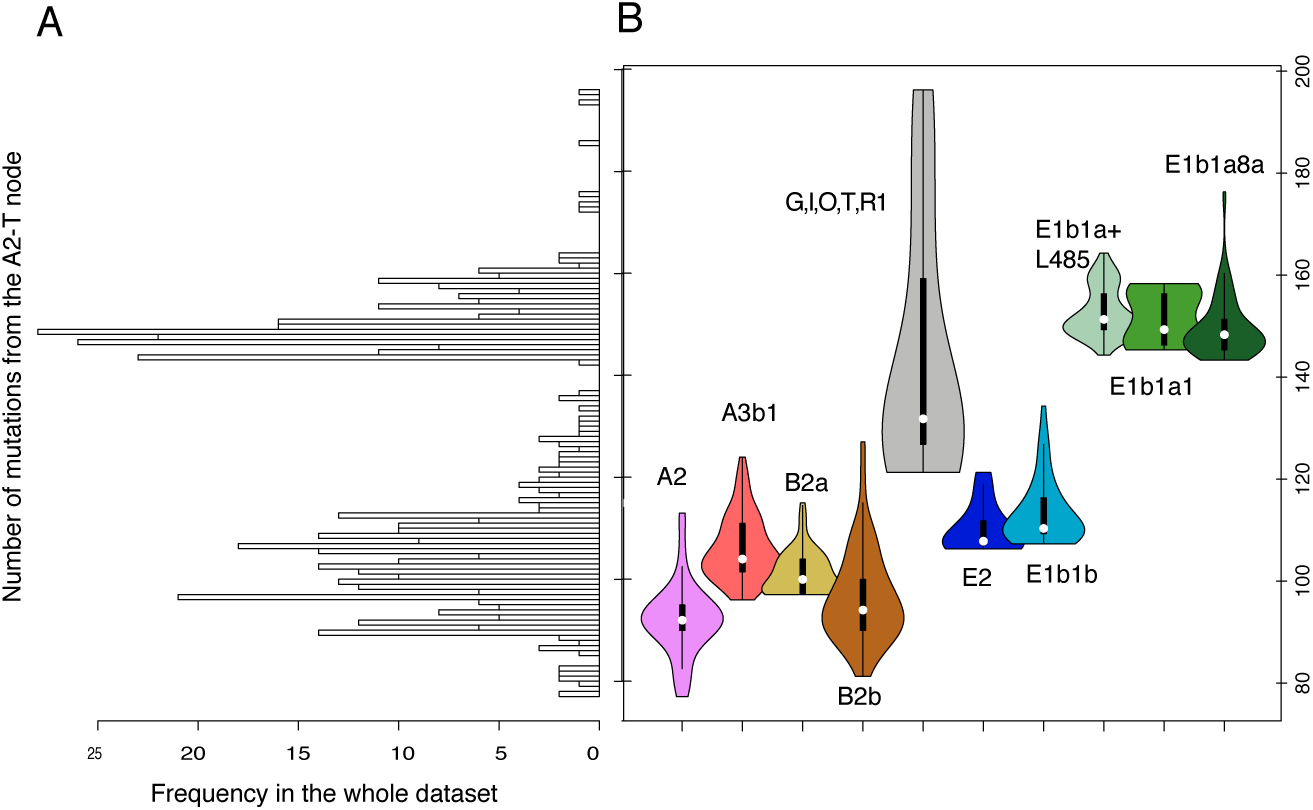
Distances to the A2-T node in number of mutations. A. Distribution of distances from each tip to the A2-T node. B. Density distribution of distances to the A2-T node for each major haplogroup. Haplogroups are color-coded as in Figure 1.

#### Impact of imputation on phylogenetic reconstruction

As described in the Material & Methods, simulations were performed to test whether the imputation process has an impact on the phylogenetic analysis and on the TMRCA dates, since imputation might reduce the observed genetic distance between sequences, which BEAST uses to reconstruct the phylogeny. When analyzing the total deviation of the node heights between trees that were constructed from alignments with different numbers of imputed sites and a tree without missing data (Supplemental Figure S12A), the results for both the strict clock and the ULN clock were approximately the same, with slightly bigger deviations for the strict clock. This probably reflects an increasing effect of removal of singletons, which by definition are located on branch tips, and thus would lead to increasing rate variation in the tips. A relaxed clock would allow for this branch rate variation and hence lead to lower deviation from the expected node heights. When 50% of the sites were imputed the amount of deviation almost doubled, which indicates that the increase in lost polymorphisms is influencing the node height estimates. In order to test if imputation affects our height estimates for nodes close to the root, we analyzed the deviation of the root node height for the same data sets (Supplemental Figure S12B). For the strict clock model, there was close to no deviation in either the mean or the median of the observed root heights from the expected values, although the 95% HPD intervals were consistently lower for the observed trees than for the expected tree. For the ULN clock model, for upper boundaries of missing data ≤ 10% both the mean and median and the 95% HPD interval of the root height of the observed trees are very close to those for the expected tree. With increasing levels of missing data, the mean and the median of the root height deviates more from the expected values, with the median providing the better fit, indicating that the 95% HPD intervals are not normally distributed. Additionally, the 95% HPD intervals widen with increasing amounts of missing data, and are twice as large as the expected intervals for an upper boundary of 50%.

We additionally performed analyses on subsets of the data consisting of 253 sequences that had less than 5% missing data before imputation (the 253L subset) as well as 10 random subsets of 253 sequences (the 253H subset). All BEAST runs performed on these subsets return mean and median root heights and 95% HPD intervals of the same order of magnitude as those obtained with the simulated data (Supplemental Figure S13A). The 253L dataset results in lower root height estimates, but subsampling our dataset cannot explain this because the 253H dataset returns even lower values. When determining the root height by counting the number of mutations to the root no difference is observed between the full and less imputed datasets. In addition, the distribution of number of mutations to the A2-T node for the 253L and 253H datasets is not strikingly different (Supplemental Figure S13B); in particular, both are strongly bimodal.

With increasing amounts of imputation the inferred clock rates over the tree deviate more strongly from the expected clock rates (Supplemental Figure S14A). Similarly, while no clear deviation of the clock rate relative to node height is detectable for upper boundaries of imputation ≤ 10%, with increasing amounts of imputation, more nodes show a strong deviation from the expected mutation rate (Supplemental Figure S14B).

Overall, the results of the simulations indicate that the major features of our results (namely, older dates for the A2-T node and various haplogroups, and branch rate heterogeneity with respect to particular haplogroups) are not an artifact of the imputation procedure, but reflect features intrinsic to the dataset.

### DISCUSSION

#### Ancient structure in B2a

Haplogroup B2a was previously associated with Bantu-speaking food-producers and populations in contact with them (Berniell-Lee et al. 2009; Batini et al. 2011), with the implication that the presence of B2a in foraging communities might indicate gene flow from food producers. Our extensive dataset of both Khoisan and Bantu-speaking groups from southern Africa allows us to address the question of the origins of B2a in more detail. Haplogroups A2, A3b1 and B2b are significantly higher in frequency in the Khoisan populations, as expected (Wood et al. 2005), while haplogroups E1b1a+L485 and E1b1a8a are significantly higher in frequency in Bantu speakers (Table 1). In contrast, B2a does not differ significantly in frequency between Bantu-speaking populations (14%) and Khoisan populations (9%, excluding the Damara, who are genetically distinct from other Khoisan groups - Pickrell et al. 2012; Barbieri et al. 2014a). The presence of both Khoisan and Bantu lineages in long separated branches suggests an early divergence of the haplogroup in the two populations (Figure 2A), and the fact that the highest frequencies of B2a are found in both southern Africa and in Cameroon (Figure 2B) – the homeland of the Bantu expansion – also makes it difficult to pinpoint an exact origin.

The network based on STR haplotypes within B2a contrasts strikingly with STR-based networks for the the Bantu-associated haplogroups E1b1a8 and E1b1a+L485: in the B2a network individuals from major geographic areas tend to cluster separately, whereas the E1b1a networks show a strong signal of recent expansion and no clear geographic or population structure (see Supplemental Figures in de Filippo et al. 2011). In sum, we find no convincing evidence that B2a was brought to southern Africa solely via the expansion of Bantu-speaking peoples, in agreement with previous studies that expressed some doubt about this association (Batini et al. 2011; Scozzari et al. 2014). Instead, the strong signal of geographic structure, older coalescence time, and high differentiation of B2a lineages in Khoisan groups all support an old presence of this haplogroup in sub-Saharan Africa. B2a might have been geographically widespread long before the expansion of speakers of Bantu languages and could thus represent an indigenous component in the Khoisan populations. Therefore, the presence of B2a in southern Africa probably represents a mix of autochthonous lineages and lineages brought by the Bantu expansion.

#### TMRCA estimates and branch length heterogeneity

One of the most notable findings of our study is that all the estimated dates from the southern African data are older than the dates estimated in previous studies: at least 38 kyrs older when counting the number of mutations to the root, and 60 kyrs older when estimated with BEAST. For example, in the southern African data the mean TMRCA for the A2-T node is 178 kyrs by counting mutations and 206 kyrs using the BEAST tree and a relaxed clock model, while Poznik et al. (2013) estimated an age of 138 kyrs for the same node and using the same mutation rate. One concern with the older ages we estimate is that they might simply reflect errors introduced by imputation. However, our results indicate that whereas estimates based on relaxed clock models and imputed data should indeed be viewed with caution (Supplemental Figure S12), the TMRCA based on mutation counts is not affected (see Supplemental Figure S13). It therefore seems that the finding in the present study of an older age for the A2-T node is not an artifact caused by imputation.

A further striking result is the branch length heterogeneity that is visible both in the MP tree (Figure 1) and in the distribution of the number of mutations from each tip to the A2-T node across different haplogroups (Figure 4). This is consistent with previous observations of branch length variation in the Y chromosome tree (Scozzari et al. 2014; Hallast et al. 2015). To try to elucidate the cause(s) of this strong branch length variation effect in the southern African data, we first investigated the potential impact of imputation with a simulated dataset, as discussed above. The results of these simulations indicate that imputation can indeed introduce rate heterogeneity, primarily by losing singletons (Supplemental Figure S3), resulting in an increasing variation of the clock rate across branches (Supplemental Figure S14). However, the simulation results also indicate that given the amount of imputation carried out on the southern African data (on average 10% missing data per individual before imputation, Supplemental Figure S15), the effect on subsequent analyses should be negligible (see Supplemental Figures S3, S12–14). Moreover, the observed branch length heterogeneity is strongly associated with particular haplogroups (Figure 4B), but there are no significant differences in the number of imputed sites across haplogroups (Supplemental Figure S16).

Branch rate variation over a phylogenetic tree can also have natural causes; in particular, population growth can cause an increase in the number of neutral mutations per chromosome (Gazave et al. 2013), and it is possible that imputation might lead to a similar signal, i.e. a tendency to a higher branch rate variation close to the tips. However, as shown in the simulated dataset, while the discrepancies in mutation rate indeed increase with increasing levels of imputation, they are distributed all over the tree. This demonstrates that imputation does not lead to the branch rate variation signal expected for an expanding population. Thus, the observed rate heterogeneity cannot be attributed to imputation.

The shortest branches in the Y chromosome phylogeny are for haplogroups A and B (Figure 1, Figure 4), and there are technical biases that could account for this. First, the array is designed with probes matching the reference genome, which is almost entirely from a haplogroup R1b individual (Xue et al. 2009); the capture could therefore favor sequences that are more similar to the reference genome. Second, the variant calling procedure in GATK is prone to accept a SNP when it is already reported as a variant in the reference genome and in a reference dataset (DePristo et al. 2011); this reference dataset is compiled from publicly available sources, in which A and B sequences are underrepresented. Third, during imputation our dataset is compared to the data from the HGDP-CEPH panel analyzed previously (Lippold et al. 2014), in which A and B sequences are similarly underrepresented. Therefore, there is less chance of imputing a variant allele at a missing position in the A and B sequences. All the biases listed above might decrease the recovery of SNPs in A and B sequences and hence contribute toward the observed branch shortening for these haplogroups.

However, these potential technical biases cannot account for all of the observed rate heterogeneity. In particular, we note that all lineages within haplogroup E are equally related to the reference genome and to non-African haplogroups, and therefore should be equally influenced by any technical bias. Nonetheless, there is marked rate heterogeneity within haplogroup E: E1b1a lineages have significantly longer branches than E1b1b or E2 lineages (W=15904, p<0.001, Figure 1, Figure 4). Sequencing coverage and/or sequence errors cannot explain the differences in branch length within haplogroup E; when removing all samples from haplogroup E that have an average coverage <10x, the branch length heterogeneity between E1b1a lineages and E1b1b and E2 lineages (Supplemental Figure S17A) is maintained (Supplemental Figure S17C, S17D). Moreover, more stringent filtering of SNPs does not eliminate the differences in branch length either (Supplemental Figure S17B, S17D).

Notably, the E1b1a lineages are all associated with Bantu-speaking populations, whereas the E1b1b and E2 lineages are not, which suggests that demographic factors associated with the Bantu expansion might be contributing to the observed rate heterogeneity. One possibility is the effect of a population expansion on the number of mutations per lineage (Gazave et al. 2013), and indeed the BSPs for the E1b1a subhaplogroups do show somewhat larger population expansions than those inferred for the other haplogroups, especially A and B (Supplemental Figure S11). Another possibility is differences in average paternal age, as suggested previously (Hallast et al. 2015), as a higher mutation rate is associated with older paternal age (Thomas 1996; Sun et al. 2012; Kong et al. 2012). For the Ju|’hoan North (known as the Dobe !Kung in previous literature), the average age of paternity is 35.8 years and the oldest documented age at last reproduction for men is 54 years (Howell 1979). In contrast, the average paternal age among agropastoralist Sub-Saharan Africans ranges from 42 years in the Herero to 46 in rural Gambians and 46.6 in Cameroon (Cochran and Harpending 2013), with the oldest age at last male reproduction in rural Gambians being 78 years (Vinicius et al. 2014). These differences in male reproductive patterns are correlated with polygyny, with the forager populations showing both the shortest span of male reproduction and the lowest levels of polygyny, whereas the longest span of male reproduction occurs in populations with the highest levels of polygyny (Vinicius et al. 2014). Mutations increase linearly with paternal age, and a 15-year increase of paternal age results in a 50% increase in mutations (Kong et al. 2012). Societal differences in average age at paternity and length of reproductive span might therefore have a considerable impact on the Y-chromosomal mutation rate over a long period (Cochran and Harpending 2013), and this might contribute to the accelerated rate of mutation we find in the Bantu-associated haplogroups, as well as the rate variation detected in other studies of human Y-chromosome variation (Hallast et al. 2015; Scozzari et al. 2014).

In conclusion, the large number of sequences from haplogroups A and B in the southern African dataset reveal new variation in these basal haplogroups and refine our understanding of the distribution of haplogroup B2a in Africa. Another important outcome is the older dates of the A2-T basal node and of individual A and B haplogroups than those published previously. In addition, we find significant rate heterogeneity in the Y chromosome phylogeny, with an accelerated rate of mutation in the Bantu-associated haplogroups. To some extent this might be attributable to the biases in SNP calling intrinsic to the method, but demographic factors, such as older average age of paternity and/or a larger population expansion in the polygynous Bantu-speaking agropastoralists, must also have contributed to the rate heterogeneity.

### METHODS

#### Sample

Individuals from Khoisan and Bantu speaking populations were sampled in Botswana, Namibia and Zambia (Pickrell et al. 2012; Barbieri et al. 2014b; Supplemental Figure S1) with the approval of the Ethics Committee of the University of Leipzig, the Research Ethics Committee of the University of Zambia, the Ministry of Youth Sport and Culture of Botswana (Research permit CYSC 1/17/2 IV (8)), and the Ministry of Health and Social Services of Namibia (Research permit Ref-Nr. 17/3/3). Each voluntary participant gave his formal consent after being told about the purpose of the study with the help of a local translator. Samples from individuals whose father and paternal grandfather belonged to the same ethnolinguistic group were selected for the study, and details on the individual samples are included in Supplemental Table S2. DNA was extracted and processed with a modified salting-out method (Quinque et al. 2006). The Damara speak a Khoisan language, but were not grouped with the other Khoisan because of their distinctive genetic background (Pickrell et al. 2012).

#### Sequencing

Bar-coded Illumina sequencing libraries prepared previously (Barbieri et al. 2013, 2014a) were enriched for ~500 kb of target NRY sequence using the Agilent Array and methods described previously (Lippold et al. 2014). Reads were generated from 7.5 lanes of the Illumina GAII (Solexa) sequencer and were mapped to hg19 with BWA (v 0.5.10 customized in-house following the guidelines in https://github.com/udo-stenzel/network-aware-bwa). In total, we generated 95,622,812 reads that passed the quality check and duplication removal and mapped to the Non-recombinant portion of the Y chromosome (NRY) region.

SNPs were called both in the target region of ~500 kb reported previously (Lippold et al. 2014) as well as in the flanking 500bp of each target region covered by the reads, giving a total of 964,809 callable sites. To improve SNP calling, the reads were merged with the available HGDP-CEPH data (Lippold et al. 2014). SNP calling and quality filtering were performed with GATK with the following settings: QD <2.0, MQ<40.0, FS>60, Haplotypescore>13.0, MQrankSum<-12.5, ReadPositionRankSum<-8, MQ0>3 and 10*MQ0>DP (as recommended by

http://gatkforums.broadinstitute.org/discussion/2806/howto-apply-hard-filters-to-a-call-set). Average coverage for all samples was 15.3X, with a minimum of 1X and a maximum of 41X. The results were stored in a VCF file containing information for each callable site of the target region; from this, a second VCF file was created that contained only the variable positions. Of the total of 622 sequences generated in the laboratory, we obtained enough data for 547 individuals so that they could be assembled and aligned before imputing the missing sites. The SNP L419, which resolves the split of haplogroups A2 and A3, was included in our callable regions but was removed by quality filters due to too much missing data. This SNP was nevertheless added to our SNP dataset to be able to properly resolve the deep-rooting structure of the phylogeny.

Haplogroup assignment was performed with an in-house script that matched our SNPs with the classification provided in ISOGG (http://www.isogg.org/tree/index.html). The haplogroup assignment was manually verified by network reconstructions and by comparing our sequence data with the sequence data for HGDP-CEPH individuals (Lippold et al. 2014) that had previously been typed for diagnostic SNPs (de Filippo et al. 2011; Shi et al. 2010). For the branches for which we did not have any SNPs that overlapped with those listed by ISOGG, a set of diagnostic positions were additionally typed by sequencing. Details concerning these SNPs and primers are available in Supplemental Table S6. The diagnostic SNPs typed in the laboratory were included only in the network reconstructions, and were excluded from the final alignment used for the remaining analyses.

When testing for the impact of read depth on branch rate heterogeneity (Supplemental Figure S17), all SNPs for which the alternative allele was not covered by at least 3 bases in at least one sample were removed. Additionally, samples whose average read depth in the target region was < 10x were also removed for this analysis.

#### Imputation

In order to minimize the impact of missing data, an imputation procedure was performed on the total dataset after assessing the accuracy of the method with a resampling procedure (Supplemental Figure S2). The imputation method applied here, modified from Lippold et al. (2014), replaces the missing SNP allele in each sequence by comparison to the three nearest sequences (based on pairwise distances over all sites). When all three nearest sequences have the same allele, the missing site is replaced by this allele, otherwise an N is kept. In order to test the overall performance of the new method, as well as the performance with increasing amounts of missing data, we constructed two datasets that lacked missing data: dataset A consists of 116 samples and 361 SNPs from the present study; dataset B consists of dataset A together with an additional 42 samples from the HGDP (a total of 158 samples and 361 SNPs). We then randomly masked 0%, 5%, 10%, 15%, 20%, 30%, 40% and 50% of site calls and performed imputation, repeating the procedure five times for each case, and calculated the fraction of N’s remaining after imputation as well as the error rate introduced by imputation. The new method outperformed the old method, and moreover the results obtained using dataset B were always better than those obtained using dataset A (Supplemental Figure S2).

The imputation was therefore applied to a dataset which included both the southern African sequences as well as the raw data for the CEPH-HGDP individuals sequenced previously (Lippold et al. 2014), which are characterized by a higher average coverage. In total, 547 sequences with an average of 293 missing sites (range 0–1775, Supplemental Figure S15) from our dataset plus 624 sequences with an average of 282 missing sites (range 0–1284) from Lippold et al. (2014) were included in the imputation procedure. After imputation, there were 2837 SNPs in the southern African sequences, of which 387 contained at least one N (average number of Ns per sequence = 1.5, range = 0 – 15). These 387 sites were removed from phylogenetic reconstruction in the network analysis.

#### STR typing

In order to compare the B2a lineages found in our southern African dataset with previously published data (Batini et al. 2011), we typed a set of 23 Y chromosome STR loci in the 55 samples belonging to haplogroup B2a using the PowerPlex^®^ Y23 System (Promega, Mannheim, Germany) with 30 amplification cycles and a final volume of 10 μl. PCR products were separated and detected with the Genetic Analyzer 3130xl (Life Technologies, Darmstadt, Germany). One microliter of the amplified sample was added to 10 μl of Hi-Di Formamide (Life Technologies, Darmstadt, Germany) which includes the CC5 ILS 500 Y23 internal length standard (Promega, Mannheim, Germany). The following electrophoresis conditions applied: polymer POP-4, 10 sec injection time, 3 kV injection voltage, 15 kV run voltage, 60°C, 1800 sec run time, Dye Set G5 (FL, JOE, TMR-ET, CXR-ET, CC5). Raw data were analyzed with the GeneMapper^®^ ID-X1.1.1. (Life Technologies, Darmstadt, Germany).

#### Phylogenetic reconstructions

A Maximum Parsimony (MP) tree (Figure 1) was generated using the Parsimony Ratchet algorithm (Nixon 1999) as implemented in the R package phangorn (Schliep 2011). The Parsimony ratchet was set up with 10 random starting trees and the most parsimonious tree was kept. The tree included an A00 sequence – the most divergent human Y-chromosomal lineage found to date (Mendez et al. 2013) – as an outgroup. The data for A00 stems from the individuals from Cameroon sequenced at high coverage for the complete NRY as reported previously (Karmin et al. 2015). Since only the sites that overlapped with our set of callable sites were retained in the alignment, these sequences were not distinct anymore; the alignment was thereby increased by 227 SNPs private to A00. The nomenclature used here follows that of Karafet et al. (Karafet et al. 2008); other nomenclatures and defining mutations are provided in Supplemental Table S1.

Network analysis was carried out to analyze the relationships among sequences and to aid in counting the number of mutations from each tip to the A2-T node. Median joining networks were calculated with Network 4.6.1.3 (Fluxus Technology, www.fluxus-engineering.com) and plotted with Network Publisher.

Network analysis was also applied to visualize relationships between the STR haplotypes determined for haplogroup B2a and including data for 16 STRs from Batini et al. (2011). In this case weights that were inversely proportional to the variance observed in our dataset (Bosch et al. 1999) were assigned to each individual STR locus. Individuals from the published dataset who had missing values for one or more loci were excluded from the analysis.

#### Dating divergence time and choice of mutation rates

The TMRCA of our southern African dataset was first estimated by multiplying the number of mutations from the A2-T node to each tip by the mutation rate (expressed as number of mutations per year), which is equivalent to the rho statistic (Jobling et al. 2013). As there is uncertainty concerning the Y chromosome mutation rate, four rates were used that are representative of the range proposed in the literature. These are: 1×10^−9^ mutations/bp/year, based on a single deep-rooting pedigree (Xue et al. 2009); 0.82×10^−9^ mutations/bp/year, based on the divergence between two lineages belonging to haplogroup Q and calibrated with archaeological dates for the entry into the Americas (Poznik et al. 2013); 0.74×10^−9^ mutations/bp/year, based on an internal calibration with two aDNA sequences (Karmin et al. 2015); and 0.62×10^−9^ mutations/bp/year, based on a conversion from the autosomal rate (Mendez et al. 2013). The four rates were adjusted for the proportion of callable sites and for the loss of polymorphic sites that contained Ns to estimate the TMRCA from the mutation counts extracted from the network in Figure S4. This resulted in one mutation every 1933 years for the rate of Xue et al., one every 2357 years for the rate of Poznik et al., one every 2612 years for the rate of Karmin et al., and one every 3118 years for the rate of Mendez et al. The rate from Poznik et al. (2013), which is in good agreement with a recent estimate from Icelandic pedigrees (Helgason et al. 2015), was chosen for displaying the main results.

#### BEAST analysis and settings

BEAST v1.8.0 (Drummond et al. 2012) was used to reconstruct the tree topology and date various nodes. The best-fitting substitution model, as chosen by jModelTest v.2.1.7 (Darriba et al. 2012), was General Time Reversible *(GTR)*. The tree model was set to *Coalescent: Bayesian Skyline* (Drummond et al. 2005) with a *piecewise-linear* skyline model. The analysis was performed using both a strict clock and an Uncorrelated Exponential Relaxed (UER) clock and a given constant rate. To ensure a proper placing of the root, the A00 sequence was forced as an outgroup. An invariant site correction was applied to adjust for the removal of all invariant sites from the alignment. Multiple runs were performed independently (strict clock: 2 runs; UER clock: 13 runs). The chain length was set to 100 million steps and parameters were logged every 5,000 steps. The resulting log and tree files were combined using BEAST’s logCombiner. A burn-in was removed (strict clock: 10%; UER clock: 30%) and the files resampled (only UER clock: every 28,000 steps). The quality of the combined runs was manually checked in Tracer v1.5 (Rambaut and Drummond 2009). The ESS value of the parameter treeModel.rootHeight, which is important for dating the nodes, was above 100 for all runs. Maximum clade credibility (MCC) trees were annotated using BEAST’s TreeAnnotator for each combination of mutation rate and clock model. The mean, median, and 95% HPD intervals of the node heights were extracted from the MCC trees and used for dating. In order to determine which clock model (strict vs. UER) was best supported by the entire dataset, marginal likelihood estimation (MLE) as implemented in BEAST (Baele et al. 2012, 2013) was executed (MLE chain length: 100,000 steps; path steps: 100). Path-sampling was performed and Bayes factors were calculated by comparing the marginal likelihood estimates of the UER clock to those of the strict clock. The marginal likelihood estimation decisively favored the UER clock over the strict clock for all four mutation rates (Bayes factors: log_10_BF > 50; decisive support following Kass and Raftery 1995).

For the analyses of single haplogroups, a simple HKY mutation model was chosen, applying the rate of Poznik et al. (2013), and a relaxed exponential model with chains of 50 million steps (70 million steps for E1b1a8a and E1b1a+L458, which have the largest number of individuals). ESS values were above 100 for all runs. An outgroup sequence was included in the runs to ensure the correct placement of the root.

#### Simulated dataset

To assess the impact of imputation on our results, a bi-allelic haploid dataset containing 100 individuals and 2,000 segregating sites was simulated following Sayres et al. (2014) using *ms* (Hudson 2002). A single tree was picked and the binary sequence data converted into a nucleotide alignment using the JC69 substitution model (Jukes and Cantor 1969). Missing sites (Ns) were randomly introduced into the sequence of each sample. Five upper boundaries for the maximum number of Ns were used: 5%, 10%, 20%, 30% and 50% of the number of segregating sites. The number of Ns for each sample was determined by drawing a number from either an exponential distribution or a uniform distribution (mean of the distribution: 50% of the upper boundary) and the missing bases were uniformly introduced over the sequence length. The exponential distribution mimics the situation where a majority of samples are of high quality, i.e. with a small number of missing bases, while a small number of samples are of low quality and have a large number of missing bases. In contrast, the uniform distribution mimics the situation where missing bases are distributed at random across sequences. The introduction of Ns was repeated 10 times for each upper boundary. Subsequently, imputation was carried out as described before, and the resulting alignments were compared to the original alignments to assess the number of wrongly assigned genotypes and shifts in the minor allele frequency distribution. These two measures represent the loss of observed genetic diversity per sample and per site, respectively. Wrongly assigned genotypes were defined as genotypes for which the consensus genotype of the three genetically closest neighbors was different from the original genotype before an N was introduced. In order to quantitatively measure the shift in the minor allele frequency, the number of singletons, doubletons, and tripletons were identified in the simulated dataset. Since the number of wrongly assigned genotypes and lost “n-tons” did not differ significantly between the exponential vs. uniform distribution of missing sites (Mann-Whitney U test: p-value 0.83 and 0.63, respectively), only the dataset constructed with a uniform sampling distribution of Ns was considered for subsequent analysis.

The impact of imputation on the phylogenetic analyses (TMRCA and rate variation) was investigated using the simulated data by comparing the differences between trees generated from imputed alignments with trees generated from an alignment with no imputed bases. Trees and TMRCAs for the simulated dataset were calculated with BEAST v1.8.0 (Drummond et al. 2012) as described above. The substitution model was set to JC69, the tree model to *Coalescent: Constant Size* to avoid potential biases of more complex models, and the clock rate was set to 1. Both a strict clock and an Uncorrelated Lognormal Relaxed (ULN) clock were tested, as the simulated dataset didn’t support the Uncorrelated Exponential Relaxed (UER) clock that was used for the Y-chromosomal dataset (ESS values < 20). For a subset of the runs the topology was fixed using a starting tree reconstructed with the non-imputed alignment in BEAST. This allowed a comparison of TMRCA and rate variation without having to adjust for different tree topologies. A correction for invariant sites was performed as described before. The chain length was 30 million steps and the burn-in was set to 30% to obtain ESS values ≥ 100. One MCC tree per run was annotated using BEAST’s TreeAnnotator and the tree files were analyzed using the R package *Phyloch* (Heibl; R Core Team 2014), with a focus on the variables node height and clock rate. MCC trees of alignments in which Ns were introduced and subsequently imputed were compared to the MCC tree of the simulated alignment without imputation. First, the total deviation of the node heights in a tree was quantified by summing up the squared deviation of each node height in the imputed tree from the corresponding node height in the tree without imputation (the expected tree), divided by the node height for the expected tree. Second, we tested if imputation affects our height estimates for nodes close to the root by analyzing the deviation of the root node height for the same data sets.

In order to verify the results of the simulations in the southern African dataset, we analyzed the performance of a subset of the total data, consisting of 253 samples that had less than 5% missing data before imputation. We compared this low-imputation subset (the 253L dataset) to the entire dataset and to 10 random subsets of 253 sequences (the 253H datasets) from the entire dataset, in order to control for any sample size effect. All datasets were processed as described above for the total dataset, and the mutation rate was set to 0.82×10^−9^ mutations/bp/year. The recovered TMRCAs from the BEAST analyses were compared to an independent estimate of TMRCA calculated from the count of mutations from the A2-T node.

Finally, we investigated the effect of imputation on the clock rate inferred in the BEAST analysis with the same simulated dataset. The non-imputed simulated data did not support a relaxed clock model over a strict clock model (see Supplemental Table S7; log10(BF) > 0). Only with increasing amounts of imputation did the data support a relaxed clock model (log10(BF) < 0). As done in the previous analysis of the node heights, the deviation (χ^2^) of the observed clock rates of the imputed datasets from the expected values of the non-imputed dataset was calculated and the corresponding 95% confidence interval was plotted (Supplemental Figure S14A).

Furthermore, the χ^2^ values of the clock rate were plotted over the node height to investigate whether imputation could also lead to a tendency to a higher branch rate variation close to the tips (Supplemental Figure S14B).

### DATA ACCESS

A .vcf file with the variants found in each sample prior to imputation, a .bed file with the callable regions, and a .fasta alignment with the imputed dataset used in the analysis, including variants private to A00 are available upon request.

## ACKNOWLEDGMENTS

We thank Udo Stenzel for assistance with the raw sequence data processing, Mario Vicente for assistance with the BEAST analysis, Monica Karmin and Toomas Kivisild for providing the A00 SNP data, and Gabriel Renaud, Greg Cochran, Henry Harpending, and Monty Slatkin for helpful discussion. This work was supported by a general grant from the Leakey Foundation and a Post-PhD grant (Nr. 8501) from the Wenner Gren Foundation (to BP), as well as by the Max Planck Society. Sample collection was partly funded by the Deutsche Forschungsgemeinschaft (as part of the European Science Foundation EUROCORES Programme EuroBABEL). BP is furthermore grateful to the LABEX ASLAN (ANR-10-LABX-0081) of Université de Lyon for its financial support within the program “Investissements d’Avenir” (ANR-11-IDEX-0007) of the French government operated by the National Research Agency (ANR).

